# APOE2 gene therapy reduces amyloid deposition, and improves markers of neuroinflammation and neurodegeneration in a mouse model of Alzheimer disease

**DOI:** 10.1101/2023.08.14.552850

**Authors:** Rosemary J Jackson, Megan S Keiser, Jonah C Meltzer, Dustin P. Fykstra, Steven E. Dierksmeier, Alexandra Melloni, Tsuneo Nakajima, Luis Tecedor, Paul T. Ranum, Ellie Carrell, YongHong Chen, David M. Holtzman, Beverly L. Davidson, Bradley T. Hyman

## Abstract

Epidemiological studies show that individuals who carry the relatively uncommon APOE ε2 allele rarely develop Alzheimer disease, and if they do they have a later age of onset, milder clinical course, and less severe neuropathological findings than others with Alzheimer disease. The contrast is especially stark in comparison to the phenotype associated with the major genetic risk factor for Alzheimer disease, APOE ε4, which has an age of onset several decades earlier, as well as a more aggressive clinical course and notably more severe neuropathological findings, especially in terms of the amount of amyloid deposition. Even one APOE ε2 allele improves phenotype, but it is uncertain if that is due to the replacement of a more toxic allele by APOE ε2, or if APOE ε2 has a protective, neuro-modulatory effect. Here, we demonstrate that brain exposure to APOE2 via a gene therapy approach which bathes the entire cortical mantle in the gene product after transduction of the ependyma, rapidly ameliorates established Aβ plaque deposition, neurodegenerative synaptic loss, and, remarkably, reduces microglial activation in an APP/PS1 mouse model despite continued expression of human APOE4. This result suggests a promising protective effect of exogenous APOE2, revealing a cell non-autonomous effect of the protein on microglial activation. We also show that plaque associated microglia in the brain of patients who inherit APOE2 similarly have less microglial reactivity to plaques. These data raise the potential that an APOE2 therapeutic could be effective in Alzheimer disease even in individuals born with the risk ε4 allele.

**One Sentence Summary:** Introduction of ApoE2 using an AAV that transduces the ependymal cells of the ventricle causes a reduction in amyloid load and plaque associated synapse loss, and reduces neuroinflammation by modulating microglial responsiveness to plaques.

## INTRODUCTION

Inheritance of the ε4 allele of apolipoprotein E (APOE) is the strongest genetic risk factor associated with the sporadic form of Alzheimer’s disease (AD), increasing risk by more than 10-fold compared to the common APOE ε3/3 genotype. By contrast, the rare APOE ε2 allele has the opposite effect ^1, 2^, and individuals with APOE ε2/2 almost never develop AD. The presence of even a single allele of APOE ε2, in individuals who are APOE ε2/3 or APOE ε2/4, have improved phenotypes compared to APOE ε3/3 or ε4/4 respectively ^2–6^. APOE ε2 carriers who do develop AD have a later age of onset, a slower rate of progression, and fewer neuropathological changes in their brains. ^5, 7, 8^. These data suggest that APOE ε2 either has a modulating effect on AD progression, or that APOE2 protein acts as a null, but “lowers the dose” of more toxic APOE3 or APOE4.

APOE isoform has been shown to effect several AD related phenotypes. APOE4 has been shown to increase the aggregation of Aβ as well as decrease the clearance of Aβ across the blood brain barrier (BBB)^9–12^. This is associated with increased AB deposition in patients with AD and this strong APOE4 and APOE2 associated phenotype is replicated in targeted replacement animal models where APOE ε4 carriers have more plaque deposition than APOE ε3 carriers which have more than APOE ε2 carriers ^9, 13^. APOE4 markedly enhances tau-related toxicity in animal models as well ^14^. APOE4 is known to be associated with synapse loss near plaques ^15, 16^ in AD patients. Finally, APOE4 has also been shown to impact neuroinflammation. APOE4 in microglia leads to a robust “proinflammatory” phenotype ^17^ although whether this is due to a cell autonomous or non-cell autonomous mechanism is unknown. Although the exact mechanism of APOE4 action in amyloid deposition, synapse loss, and neuroinflammation is unknown, it appears that, in both humans and murine models, inherited APOE2 prevents or delays these features ^7, 8^.

APOE is a secreted protein predominantly expressed by astrocytes and microglia in the central nervous system (CNS); peripherally it is largely made in the liver. The BBB precludes APOE transport in both directions, so that CNS APOE reflects only protein manufactured within the CNS ^18^. As a step towards testing if APOE2 protein delivered to the CNS might be beneficial for patients with APOE4 related Alzheimer disease, we developed a gene therapy approach. Since Alzheimer’s disease impacts the entire cortical mantle, we refined a method to express APOE2 from the ependymal lining of the ventricles, allowing for secretion of APOE2 into the cerebrospinal fluid (CSF), neuropil, and interstitial fluid. The method involves use of an ependymal specific promoter to restrict expression to ependymal cells. We demonstrate that APOE2 secretion using this method ameliorates the Aβ, neuroinflammatory and neurodegenerative phenotypes seen in a widely used APPPS1 mouse model of AD, here having been crossed with human APOE4 targeted replacement alleles to generate an APOE4/4 model of Alzheimer pathology^19, 20^. These data allowed us to test the hypothesis that expression of APOE2 ameliorates the amyloid, neurodegenerative, and neuroinflammatory phenotypes present in this animal model, even in the presence of APOE4 from birth. This study tests ttwo alternative models: that APOE2 has an gain-of-function modulatory role in the development and progression of AD pathology or that the benefits of inheriting and APOE2 allele are due to decreasing the amount of more toxic APOE3 or APOE4. Since our experimental approach was to add APOE2 to a stable amount of APOE4, the data clearly favor a modulatory effect of APOE2 expression in AD related pathological changes. Moreover the effect of APOE impact on microglia appears to be non-cell autonomous. We further evaluated whether the presence of APOE2 had a similar effect on microglial phenotype in human AD tissue and found that plaque associated microglia in APOE2 carriers have less active characteristics. Together, these data suggest that introduction of APOE2, even without changing APOE4, may be beneficial in patients with the most aggressive common form of late onset Alzheimer disease.

## RESULTS

### Intraventricular injection of AAV.APOE2 leads to sustained production of APOE2 in the brain in a dose dependent manner

We performed a single intracerebroventriclar (ICV) injection of AAV causing ependymal expression of APOE2^21^ in four-month-old *Apoe* KO mice, which were sacrificed 2 months later. Injection with 7E^10^ viral genomes (vg) of virus into *Apoe* KO mice showed robust expression of APOE2 mRNA in the ependymal cell lining as detected using RNAscope for human APOE (Figure 1A). APOE2 protein was also detected in a TBS extraction of the cortex assessed by western blots and is about 10 % of APOE present in the APOE4 target replacement mice (Figure 1B,C).

**Figure1.**
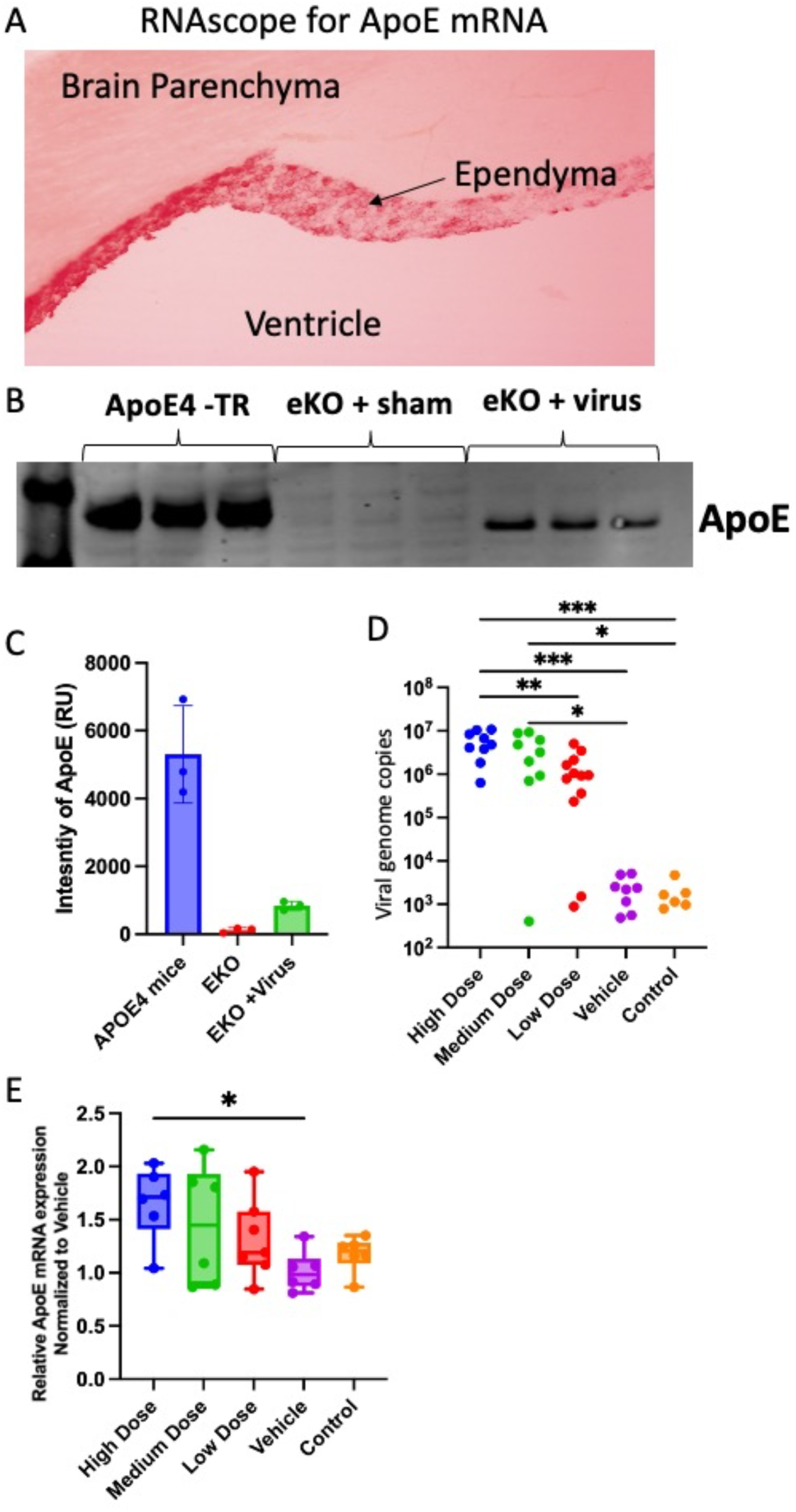
Ependymal cell expression of APOE2 driven by a AAV.APOE2. **(A)** In situ hybridization showing human APOE expression in the ependymal cells of the ventricle in a *Apoe* KO mouse. **(B)** Western blot for APOE showing that ependymal-expressed APOE2 in the cortex of the *Apoe* KO mice is approximately 10% **(C)** that of endogenous level. **(D)** Viral genome copies in tissue extracted from each mouse [(F (4, 35) = 5.546 p=0.0014) Post Hoc Tukey’s multiple comparisons test]. **(E)** qRTPCR for human APOE relative to mouse β-actin and normalized to vehicle treated animals (F (4, 26) = 2.890 p=0.0419 Post Hoc Dunnett’s test to vehicle). (E) Line graph showing a significant correlation (p=0.0445) between mRNA for APOE and viral genome copies. N indicated as each mouse is an individual dot. * p<0.05, ** p<0.01, ***p<0.001.

The AAV.APOE2 was injected ICV at three different doses: Low-7E^9^ vg, Mid-2E^10^ vg, and High-7E^10^ vg, as well as a vehicle control group into four-month-old APPPS1/APOE4 animals. Two months later, DNA extraction from the hindbrain followed by qPCR against the promotor region of the AAV plasmid showed a dose dependent effect on uptake; three animals showed no expression and were dropped from the study (Figure 1D). The number of viral genome copies correlates with increased human APOE mRNA levels which is ∼50% higher in high dose animals than baseline (Figure 1E). The transgenic APOE2 diffuses throughout the cortex, as shown by its association with plaques throughout the cortical mantle, which can be visualized immunostained with an APOE2 specific antibody (Supplemental Figure 1).

### Expression of APOE2 has a dose dependent effect on the level of AB plaque deposition

At four months of age (time of injection) APPPS1/APOE4 animals show modest plaque deposition and by six months of age (time of necropsy) plaque deposition is well established across the cortex (Supplemental Figure 2). Using ThioS as a marker for dense core amyloid plaques we show an effect of APOE2 leading to reduced plaque deposition (Figure 2 A-C). The high dose animals show a significant ∼33% reduction in the percent of the cortex covered by ThioS positive staining as compared with the vehicle treated animals (Figure 2B). The dose dependent effects of APOE2 on plaque burden correlates with the amount of DNA expression seen in each individual animal (Figure 2C). Staining using an anti-Aβ (Aβ) antibody showed a similar trend at both the group (Supplemental Figure 3A) and the individual level (Supplemental Figure 3B).

**Figure 2:**
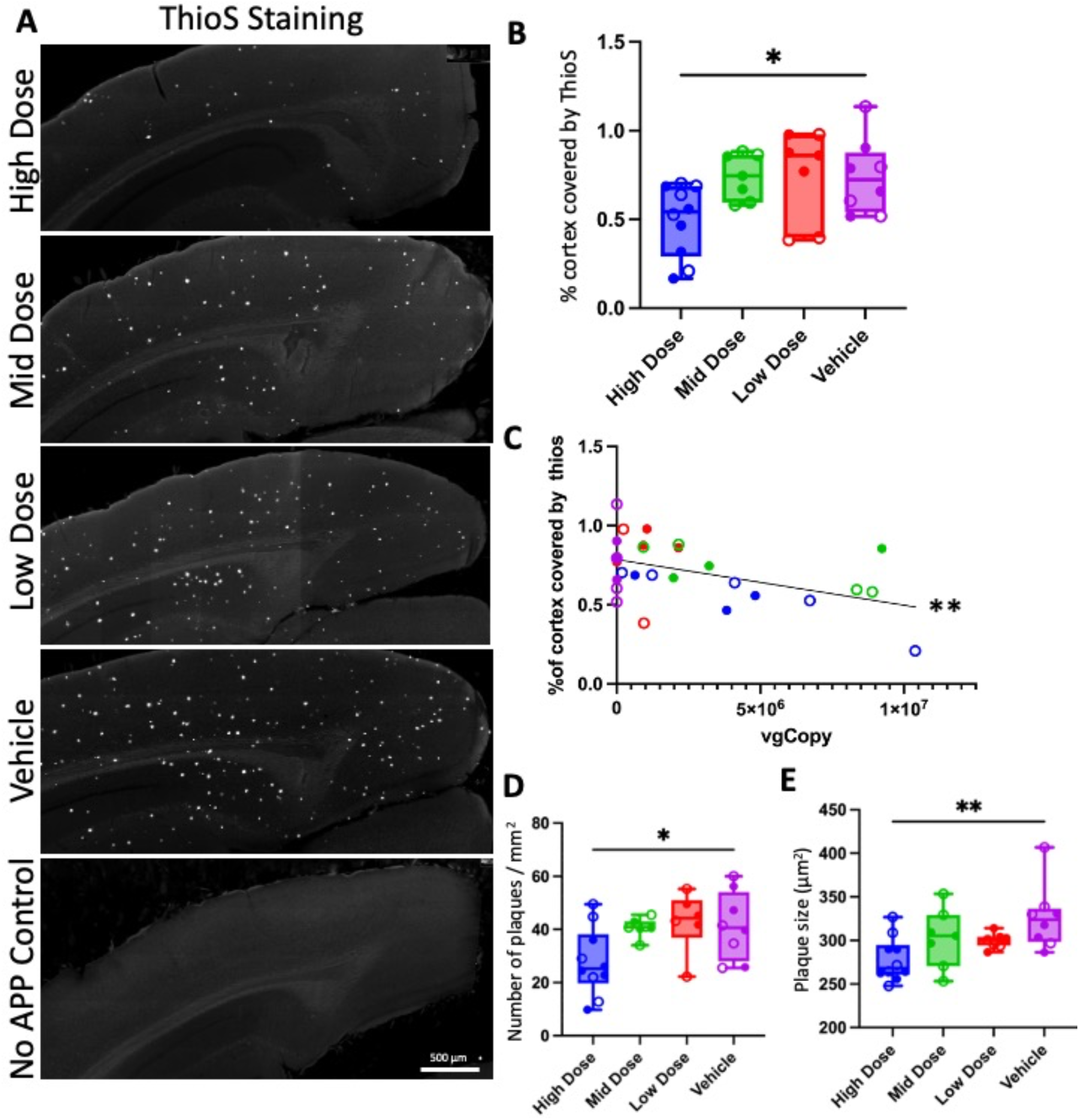
APOE2 reduces plaque deposition, number, and size in a dose dependent manner. **(A)** IHC for ThioS in the cortex of dosed APPPS1/APOE4 animals. **(B)** Percent cortex coverage by ThioS staining is significantly lower in the high dose animals (F (3, 27) = 4.310 p=0.0329). **(C)** The percent cortical coverage by ThioS is significantly correlated (p=0.01) to the number of viral genome copies in the brain sample from each mouse. **(D)** Plaque number (F (3, 27) = 3.597 p=0.0263) and **(E)** plaque size (F (3, 27) = 4.113 p=0.0159) both show a significant effect in the high dose group. n indicated as each mouse is an individual dot with open circles as females and closed as males Post Hoc tests are shown as Dunnett’s multiple comparisons test comparing with vehicle. p * p<0.05, ** p<0.01.

**Figure 3:**
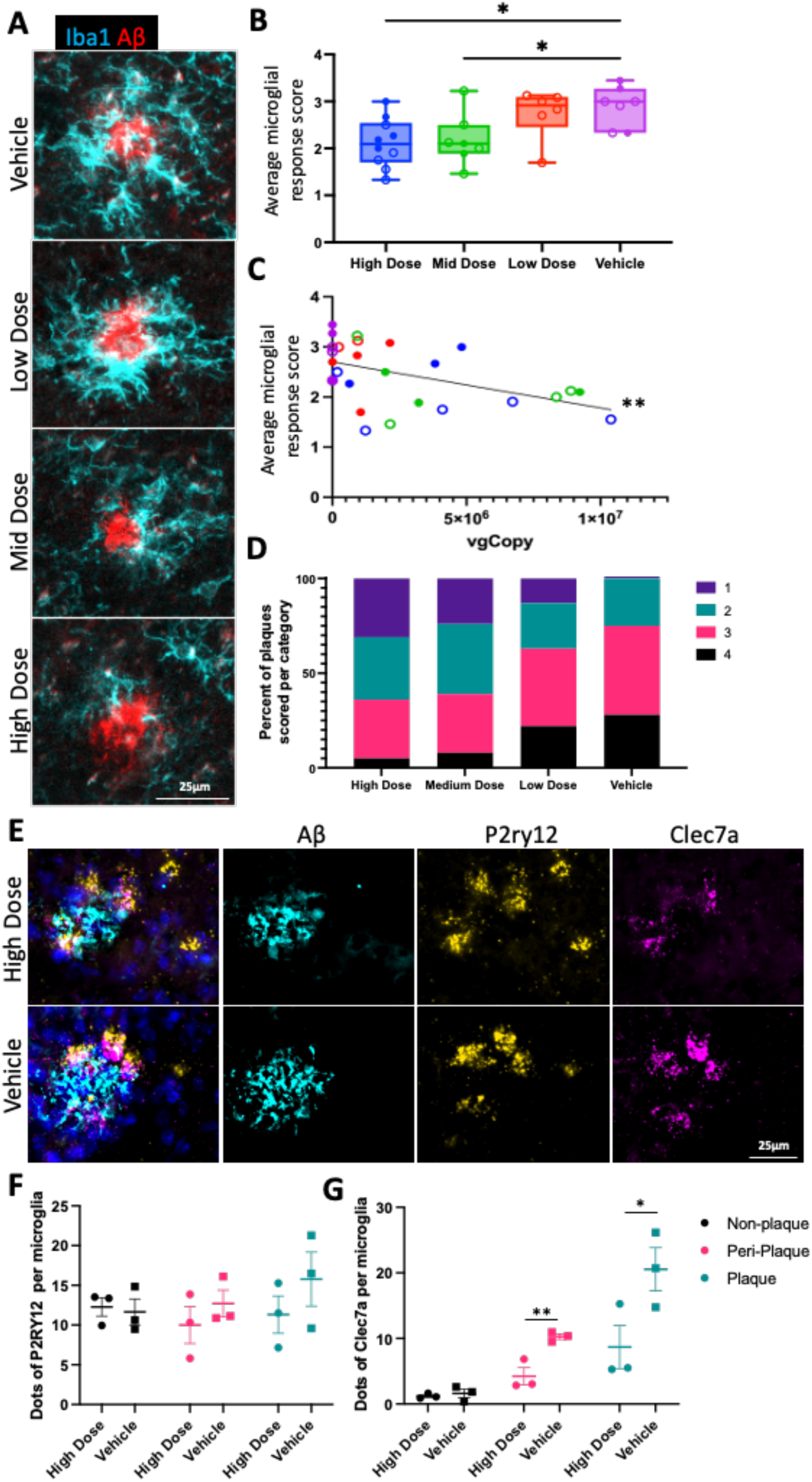
APOE2 reduces microgliosis near plaques. **(A)** IHC for IBA1 and Aβ in the cortex of dosed APPPS1/APOE4 animals. **(B)** average microglial response score (F (3, 26) = 4.529 p=0.0110) shows a reduction in microglial activation in the high and mid dose groups and **(C)** is significantly correlated (p = 0.0081) to the number of viral genome copies in that mouse. **(D)** This reduction is due to a decrease in the number of plaques scored as a 4 and an increase in the number of plaques scored a 1. n indicated as each mouse is an individual dot with open circles as females and closed as males. Post Hoc tests are shown as Dunnett’s multiple comparisons test comparing with vehicle. **(E)** RNAscope for P2ry12 and Clec7a of high dose and vehicle treated animals shows that there is **(F)** no change in the number of P2ry12 transcripts per microglia in the treated animals there is a **(G)** significant attenuation in the number of Clec7a transcripts per microglia in the plaque (p=0.0317) and the peri-plaque area (p=0.0057) in the high dose animals compared with controls p * p<0.05 ** p<0.01.

This reduction corresponds to both a significant reduction in plaque density (Figure 2D) and an even more significant reduction in the size of Aβ plaques (Figure 2E) when high dose animals are compared with vehicle treated animals.

Biochemical measures of amyloid align with, and confirm, the pathological measures. The concentrations of Aβ42 peptides measured from the formic acid and SDS soluble extracts of mouse brain mimicked the changes observed histologically such that the high dose animals showed a ∼50% reduction in the amount of both SDS (Supplemental Figure 3C) and formic acid (Supplemental Figure 3D) soluble Aβ42.

### Expression of APOE2 has a dose dependent effect on plaque related neuroinflammation

Data from human patients show that microglia in APOE ε4 patients have a more inflammatory phenotype than APOE ε3 or 2 individuals ^17^. To see if the addition of APOE2 could attenuate the effect of plaques on plaque associated glia we performed IHC for Aβ, Iba1 (microglia) (Figure 3A, Supplemental Figure 4A) and GFAP (astrocytes) (Supplemental Figure 4A,B). We assessed the level of plaque associated glial reactivity on a semiquantitative 4-point scale where 1 is non-reactive and 4 is very reactive, assessing the areas immediately surrounding the plaque (Supplemental Figure 4A). The scale was validated by having the images assessed by 2 independent investigators (RJJ and BTH), with a correlation R^2^ of 0.89.

**Figure 4:**
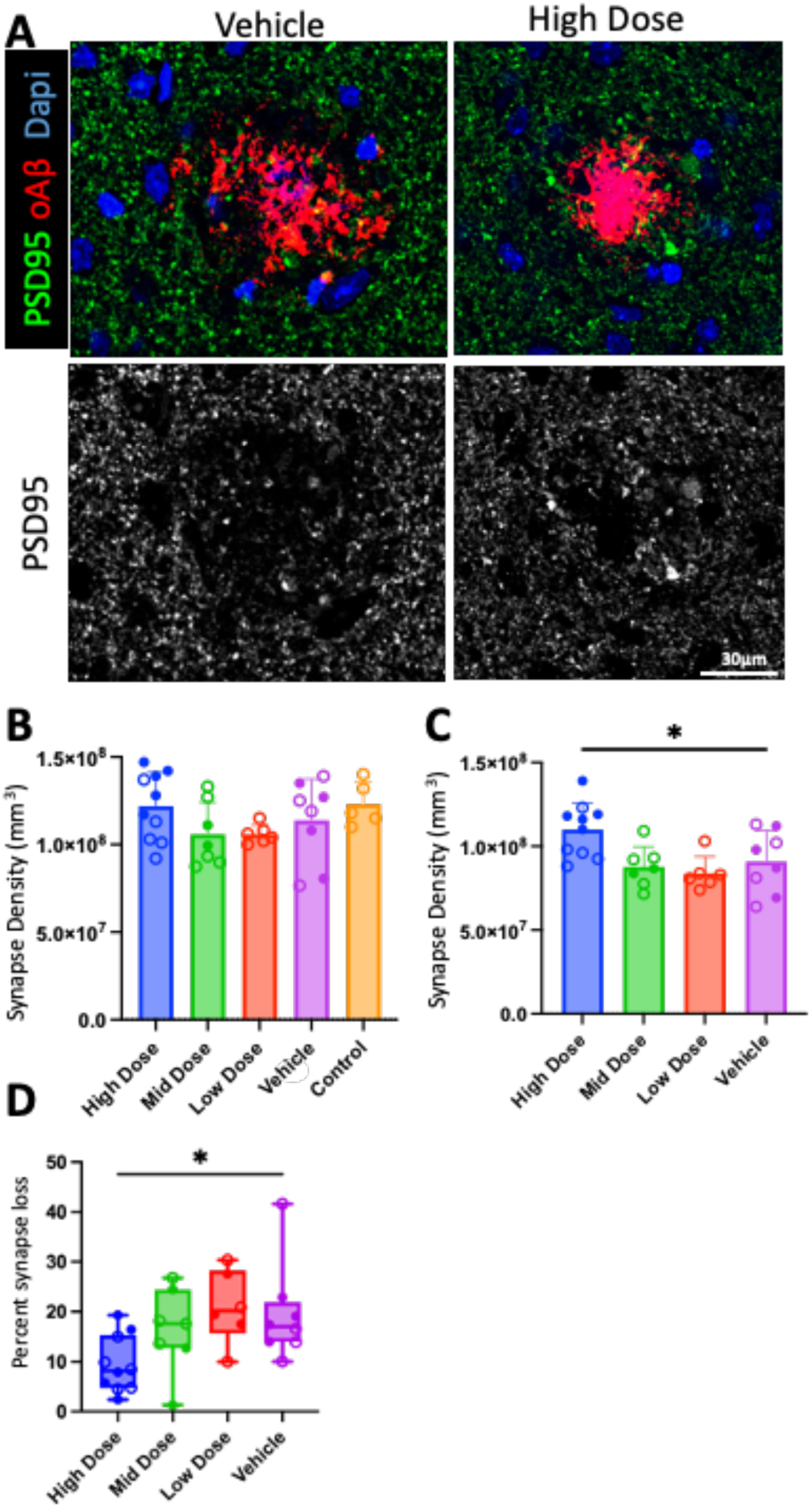
APOE2 reduces synaptic loss near plaques. **(A)** IHC for PSD95 and Aβ in the cortex of dosed APPPS1/APOE4 animals. **(B)** Synapse density is unchanged far from plaques but **(C)** significantly increased near plaques (F (3, 27) = 5.153 p=0.0060). **(D)** this leads to a significant decrease in the percent synapse loss in the high dose animals compared with vehicle (F (3, 27) = 3.693 p =0.0239). n indicated as each mouse is an individual dot with open circles as females and closed as males. Post Hoc tests are shown as Dunnett’s multiple comparisons test comparing with vehicle. p * p<0.05.

We found that there is an obvious, statistically significant attenuation of microglial reactivity in the high and mid dose mice as compared with the vehicle treated mice (Figure 3B,C) and that this attenuation shows significant correlation with dose (Figure 3C). This attenuation appears to be driven by an increase in the number of plaques that are not associated with a strong microglial reaction (score of 1) in the high and mid dose animals and a corresponding reduction in plaques that have a score of 4 as compared with vehicle treated animals where almost all the plaques are associated with highly reactive microglia (Figure 3D). We further assessed the local activation state of microglia by utilizing RNAscope probes for the homeostatic microglial marker P2ry12 and the damage-associated microglial marker Clec7a ^22^ in 3 High Dose and 3 Vehicle treated animals (Figure 3E). P2ry12 puncta are found in cells that are reasonably evenly spread throughout the cortex although there are microglial clusters of cells near plaques while Clec7a mRNA is found predominantly in cells in the immediate vicinity of plaques. We assessed 50 plaques and 50 non plaque regions per mouse as well as the peri-plaque area, here defined as the region <25µm from the edge of the plaque halo, and we see no change in the amount of P2ry12 puncta per microglia when comparing plaque proximity or treatment group (Figure 3F). These data show a significant difference between treatment groups in the number of Clec7a transcripts in both the peri-plaque and plaque areas (Figure 3G) confirming the morphological data (Figure 3C) with a molecular marker of disease associated microglial activation ^23^.

Meanwhile astrocyte reactivity around plaques is unaffected by APOE2 levels or expression (Supplemental Figure 4 C,D,E) and is equally elevated around plaques in all groups.

### APOE2 exposure improves measures of synaptic loss around amyloid deposits

Synapse loss is known to correlate with cognitive impairment and has been shown to occur near plaques in human patients ^16^ as well as in this mouse model ^24^,with higher amount of synaptic loss near plaques in APOE4 compared with APOE3 mice or carriers.

Post-synaptic densities (PSD95) were stained using immunohistochemistry and imaged using a confocal microscope (Figure 4A). As synapse loss in this model occurs within ∼15 µm of the edge of a plaque ^24^. We measured synapse density within the halo of expected synapse loss (within 15µm of the plaque edge) as well as in an area far from plaques where synapse density would be expected to be normal (>40 µm from the plaque edge). As expected, synapse density far from plaques did not differ among groups (Figure 4B). Excitingly, the high dose animals showed an increased level of synapses near plaques compared with vehicle treated animals (Figure 4C), restoring this marker of neurodegeneration to near normal levels. Percent synapse loss was calculated by comparing the synaptic density near plaques with synaptic density far from plaques within the same animal (Figure 4D). Vehicle treated animals show twice as much synapse loss as high dose animals with 70% of high dose animals showing less than 10% loss near plaques as compared with the other groups where all but one mid dose animal showed more than 10% loss.

We also evaluated the number of dystrophic neurites associated with amyloid deposits by staining for the axonal marker SMI312 alongside an Aβ antibody (Supplemental Figure 5A). In this model, these dystrophic neurites generally do not immunostain for abnormal tau epitopes, and there is not a strong tau phenotype. We found no difference in the number of axonal neuritic dystrophies among the groups (Supplemental Figure 5B,C).

**Figure 5:**
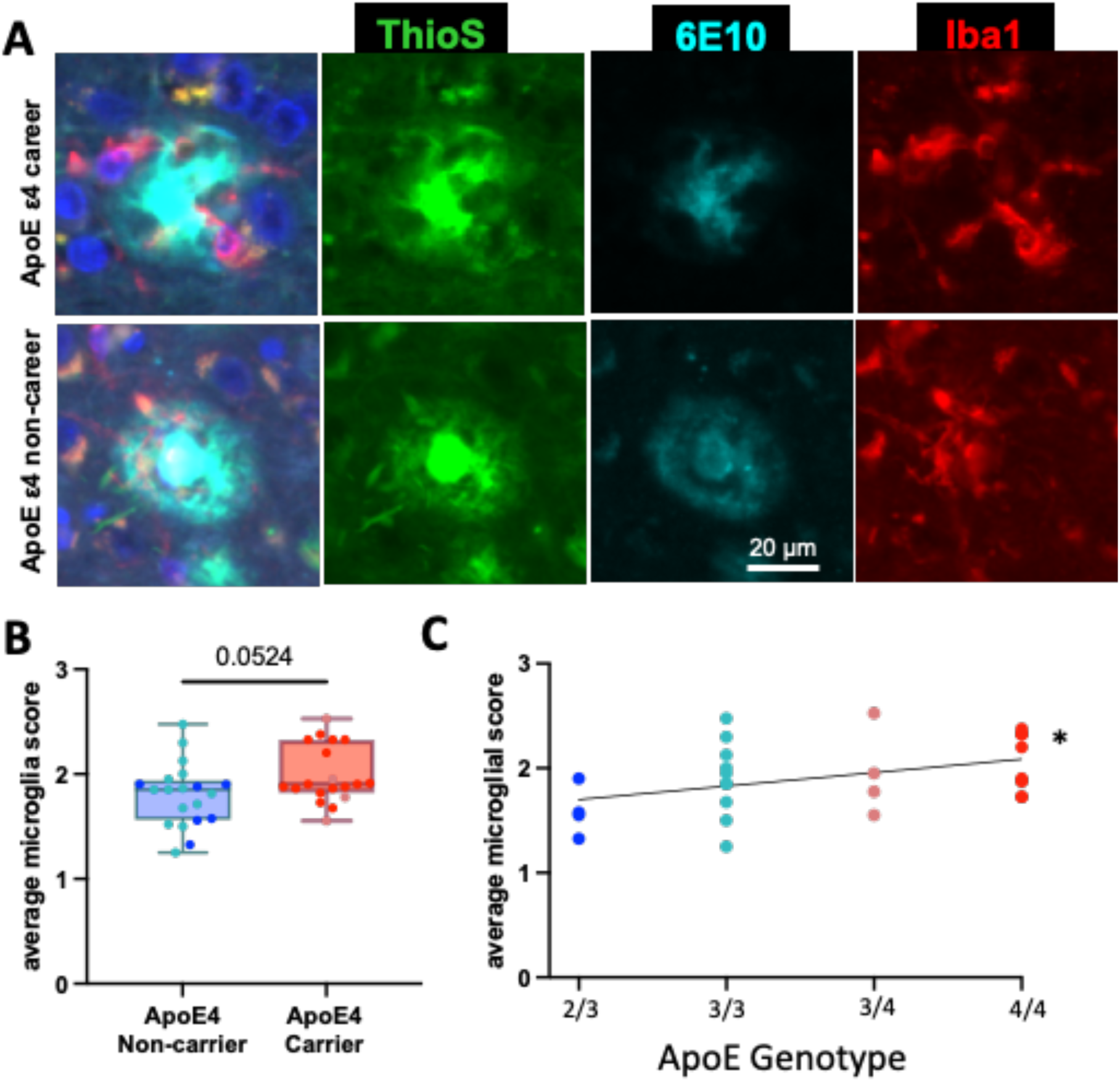
APOE2 reduces microgliosis near plaques. **A)** IHC for IBA1 and Aβ in the frontal cortex of post-mortem tissue from end stage human AD cases. **(B)** Average microglial response score (t(37)=2.0, p=0.0524) shows an increase in microglial activation in APOE4 carriers compared with non-carriers and **(C)** there is a significant correlation between microglia activation and APOE risk (where APOE2/3 individuals are assessed as have -1 APOE4 allele) (F(1, 35)=5.410 p=0.0259)

### APOE isoform has an effect on plaque related neuroinflammation in humans

To assess the translatability of our findings we looked at microglia in the immediate vicinity of plaques in frontal cortex collected from individuals with a neuropathological diagnosis of AD and known ApoE genotypes (Supplemental table 1, Figure 5A). We again assessed the level of plaque associated microglial reactivity on a semiquantitative 4-point scale where 1 is non-reactive and 4 is very reactive, assessing the areas immediately surrounding the plaque. Overall plaques in human tissue show a lower level of microglial reactivity than those in mice (Figure 3B, 5B) potentially due to the speed of plaque formation or the amount of soluble Aβ produced in mice with familial AD mutations. There was a significant correlation of microglial reactivity with the number of APOE alleles (where APOE2/3 individuals were considered to have -1 APOE4 alleles) (Figure 5C). This translated into a trend towards an increased level of microglial reactivity in APOE4 carriers were compared with non-carriers (Figure 5B).

## DISCUSSION

The strong genetic and experimental link between APOE genotype and AD has long made APOE a subject of interest when considering AD risk modifiers or therapeutics ^5, 7^. While APOE4 is unequivocally associated with a more aggressive AD phenotype, the protective effect of APOE2 has been attributed to it acting as a null, reducing the dose of the more toxic forms of APOE. For example, APOE2 protein has far less binding to the LDL receptor than other isoforms, although this is not true of binding to other members of the LDL receptor family ^25^. In contrast, we hypothesized that APOE,2 might have a positive modulatory impact on neurodegenerative processes, leading to a beneficial gain of function mechanism. Our data suggest strongly that the latter is the case, since expression of a relatively small amount (∼10% of basal amounts) of APOE2, in the setting of continued unchanged amounts of APOE4, strongly impact AD related phenotypes. The relatively small amount of APOE2 present suggests strongly that the effect is a therapeutic beneficial gain of function, rather than competition with APOE4. Importantly, in either case, this improvement is observed in the setting of mild but already established plaque deposition and continued APOE4 expression, showing that the effect of APOE2 actively modulates the impact of APOE4, rather than simply acting as a less toxic form or physiologically null allele.

The practical issues of distributing gene product throughout the brain is a barrier to using gene therapy in widely distributed diseases like AD. This is especially true in larger organisms such as non-human primates or humans where the diffusion rate of injected material through the brain parenchyma limits the ability of a single AAV injection to effect large areas of brain. Our current and previous studies suggests an approach to overcome this barrier: expression of secreted proteins via transduction of the ependyma and related structures, allowing secreted protein to diffuse throughout the cortical mantle ^24, 26, 27^. The diffusion rate of AAV through the liquid CSF held in the ventricle is much greater than that of the relatively solid parenchyma and the ependymal cell layer of the ventricle is important for the maintenance and production of the CSF^28^. Thus, transfection of these cells allows for gene product to be pumped out into the CSF where it can affect the entire cortical mantel and makes for an attractive method of overcoming the limitations introduced by a larger target area in the jump from rodent models to humans.

We applied this approach to expressing APOE2 in an APPPS1/APOE4 model of Alzheimer pathology, and show that, in achievable doses, expression of APOE2 secretion into the neuropil can positively impact plaque deposition, neuroinflammation, and neurodegeneration within a short time interval-8 weeks. The effect of APOE2 on plaques is expected from a robust neuropathological literature showing that APOE,2 carriers have fewer plaques. The results here suggests that the impact of APOE in AD is ongoing, and that manipulating APOE even after plaques are established can alter the course of the disease. APOE2 has been shown to increase the clearance of Aβ across the BBB and well as a decrease aggregation as previous studies have shown APOE effects for both the clearance and aggregation of Aβ ^7, 9, 12^. The positive APOE2 effect we see is on both dense core fibrillar plaques and on biochemical measures of Aβ, consistent with the interpretation that APOE2 is actively facilitating increased clearance of Aβ across the BBB.

Neuroinflammation and AD have been linked in part due to the number of microglial genes that are risk factors for AD ^29^ and also due to the marked increase in micro- and astrogliosis in brains from both AD cases and mouse models ^30, 31^. Microglia produce APOE under basal conditions but production is dramatically increased in these cells in the context of AD, leading to the hypothesis that microglial produced APOE4 has a toxic cell autonomous gain of function effect^23^. By contrast with this expectation, our current data show that exogenous APOE2 has a dampening effect on microglial activation near plaques in these mice (Figure 3), consistent with an additional non-cell autonomous effect. The effect of APOE2 on microglia in this paper is reminiscent of the depressed microglial response seen in TREM2 KO mice ^32^ but here we do not see a concurrent increase in neuritic dystrophies or synaptic loss that are seen in those models ^33^.

Importantly this effect of APOE on microglial reactivity to plaques is also seen in human AD cases despite the heterogeneity and relatively end stage nature inherent in studying human brain tissue (Figure 5). Here we showed a modest increase in microglial reactivity score when comparing APOE4 carriers to non-carriers. This increase occurs step wise with relative APOE risk as a test for linear trend is significant (Figure 5). Importantly this trend only became evident after incorporating the APOE2/3 individuals indicating that the presence of APOE2 might be a greater driving force in reducing microglial responsiveness near plaques than reducing APOE4.

APOE4 has been associated with more severe synapse loss near plaques in AD in both human and mouse models ^16, 24^. In this model we also see a loss of synapses near plaques however this synapse loss is lessened in the mice expressing APOE2. This could be due to reduced levels of synaptic pruning due to less active microglia, or to reduced bioactive oAβ. Alternatively, or in addition, we have previously shown that APOE and oligomeric Aβ colocalize at the synapse, and we postulated that the interaction of oAβ with APOE2 rather than APOE4 could ameliorate synaptotoxic effects. Whether the interaction of APOE2 with oAβ, or the reduction in synaptotoxic phenotype of activated microglia, or both, improve the synapse loss around plaques, the observation that APOE2 expression can improve a core feature of neurodegeneration suggests a potential role for this approach in a therapeutic context.

Reduction of Aβ following immunotherapy ^34, 35^, which leads to profound plaque reduction and modest cognitive gains, has been recognized by FDA approvals. In contrast with Aβ immunotherapy, APOE2, while also impacting plaques, does not rely on activation of microglia and increased neuroinflammation to modulate plaques, and indeed appears to dramatically quench existing inflammatory microglial signatures and restore synaptic integrity. Taken together, these results suggest that APOE2 is a promising approach for potential therapeutic development in AD and related disorders.

**Supplemental Figure 1:**
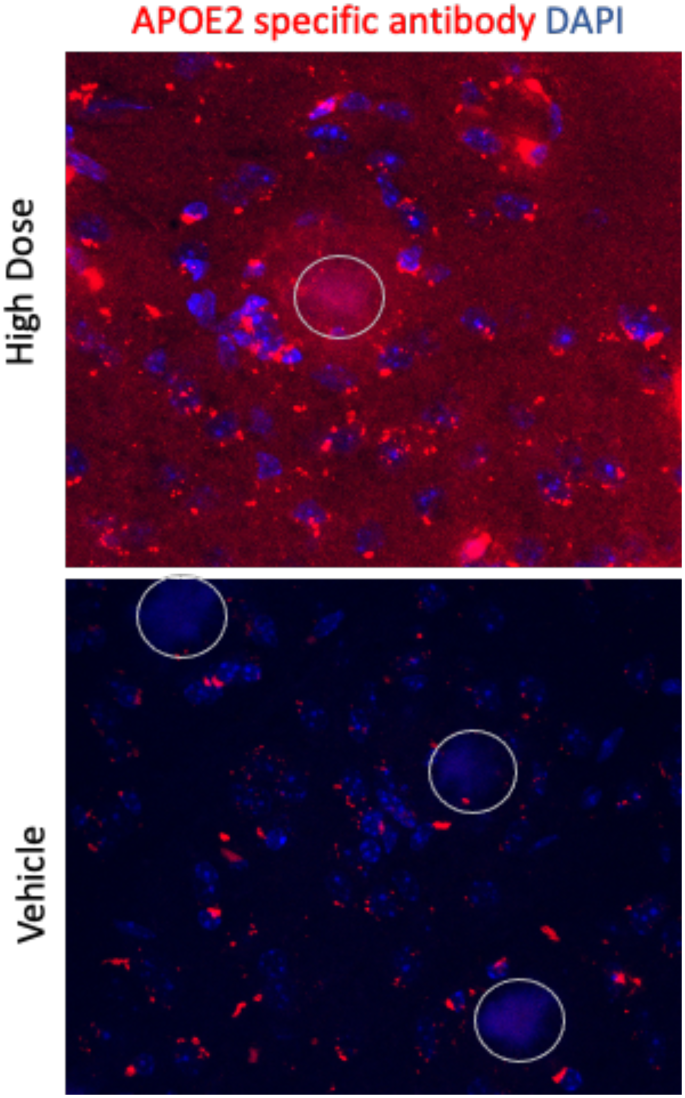
APOE2 is found throughout the cortical mantel. IHC APOE2 using an APOE2 specific antibody found diffuse staining throughout the cortex with more intense staining in the plaque (indicated by a white circle) and plaque halo.

**Supplemental Figure 2:**
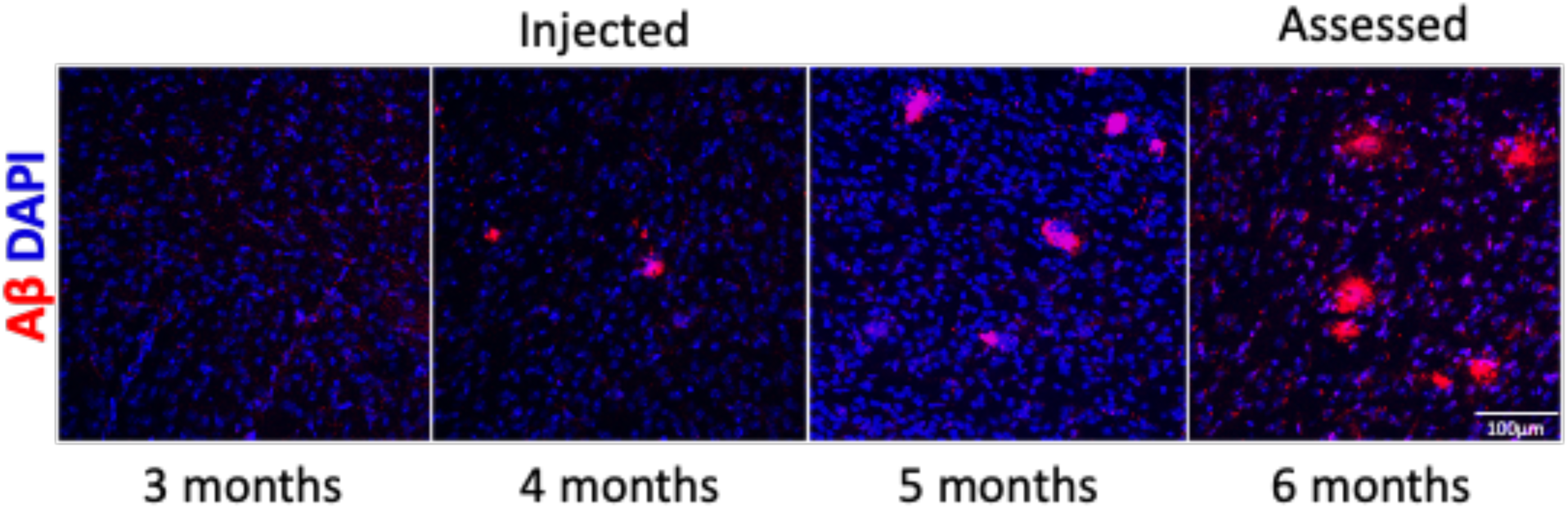
Plaque deposition in the APPPS1/APOE4 mice. Representative images of plaque deposition in the cortex in naïve APPPS1/APOE4 mice at time points that represent before, at, and after the point of injection.

**Supplemental Figure 3:**
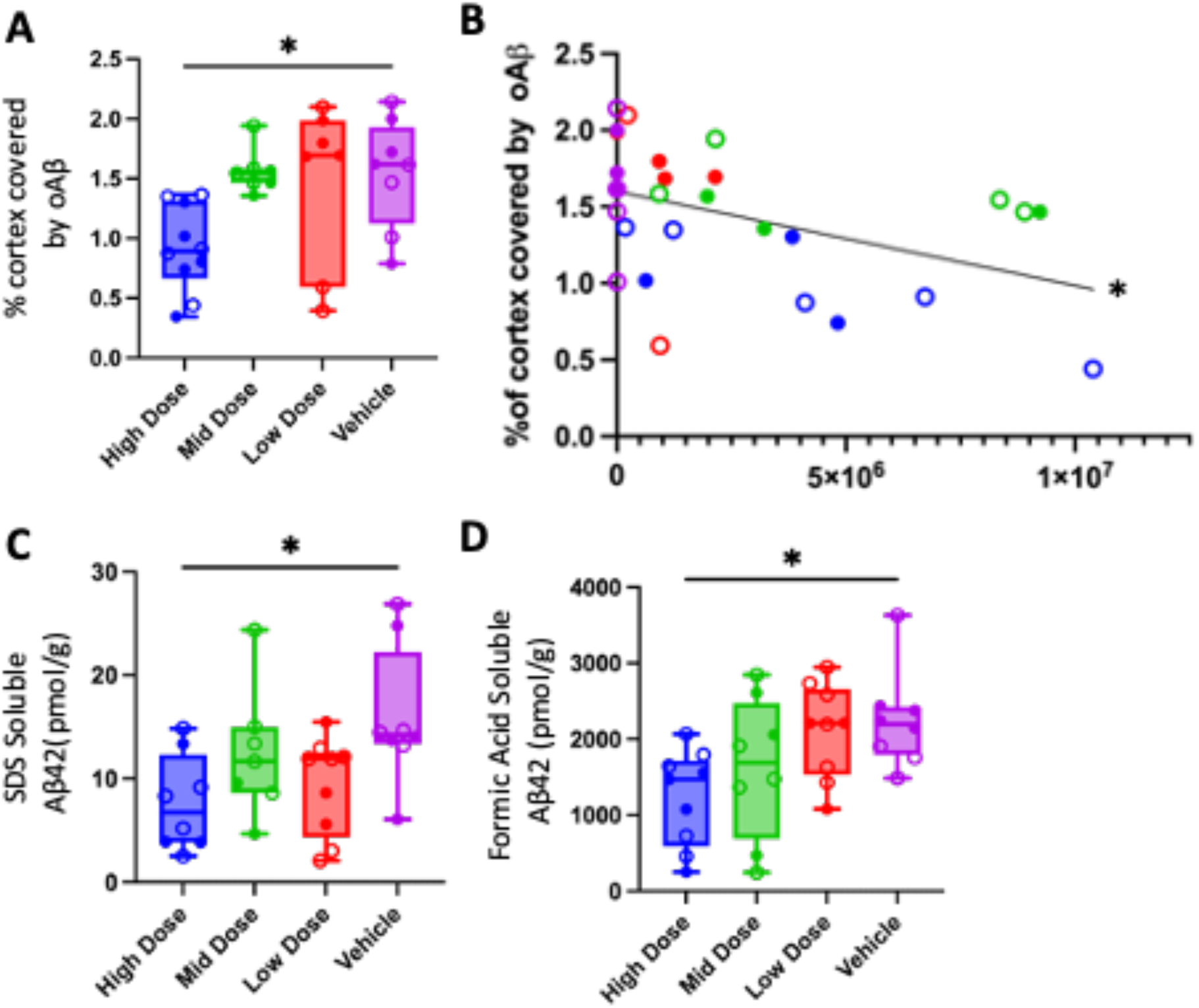
Effect of APOE2 on oligomeric Aβ. **(A)** Percent cortex coverage by Aβ staining is significantly lower in the high dose animals (F (3, 27) = 6.336, p=0.0022). **(B)** The percent cortical coverage by ThioS is significantly correlated (p= 0.0173) to the number of viral genome copies in that mouse. **(C)** ELISA shows that SDS soluble (F (3, 28) = 3.497 p=0.0285) and **(D)** Formic acid soluble (F (3, 30) = 3.741 p=0.0214) Aβ both show a significant effect in the high dose group. n indicated as each mouse is an individual dot with open circles as females and closed as males. Post Hoc tests are shown as Dunnett’s multiple comparisons test comparing with vehicle. p * p<0.05, ** p<0.01.

**Supplemental Figure 4:**
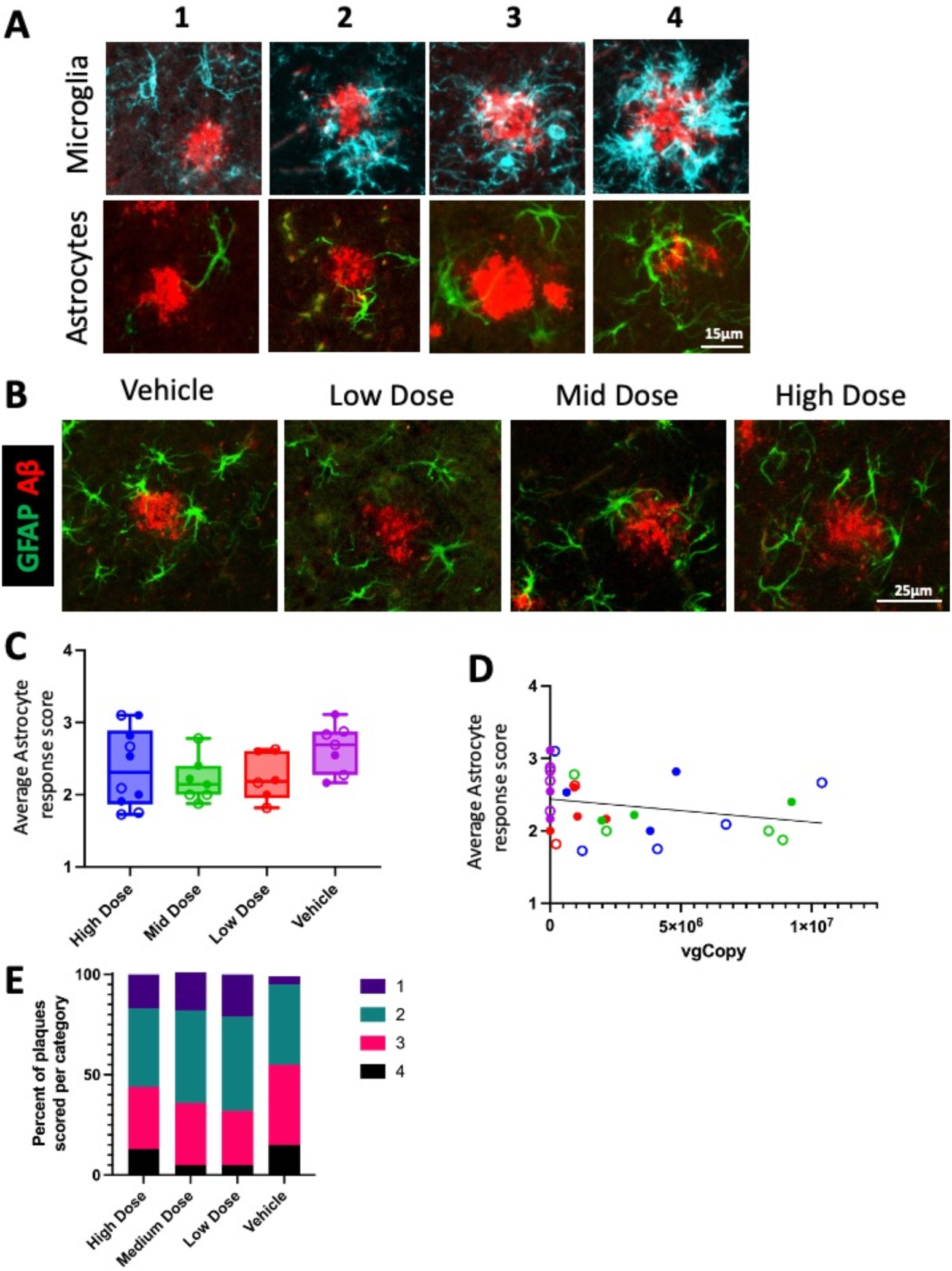
APOE2 does not affect astrocyte reactivity near plaques. **(A)** Representative images of the four point scale used to assess micro and astroglia reactivity to plaques. **(B)** IHC for GFAP and Aβ in the cortex of dosed APPPS1/APOE4 animals. **(C)** Average astrocyte response score (F (3, 26) = 1.634 p=0.2057) shows no change between groups and **(D)** is not significantly (p=0.1904) correlated to viral genome number. **(D)** There is no difference between groups in the number of plaques scored into each category. n indicated as each mouse is an individual dot with open circles as females and closed as males. Post Hoc tests are shown as Dunnett’s multiple comparisons test comparing with vehicle. p * p<0.05.

**Supplemental Figure 5:**
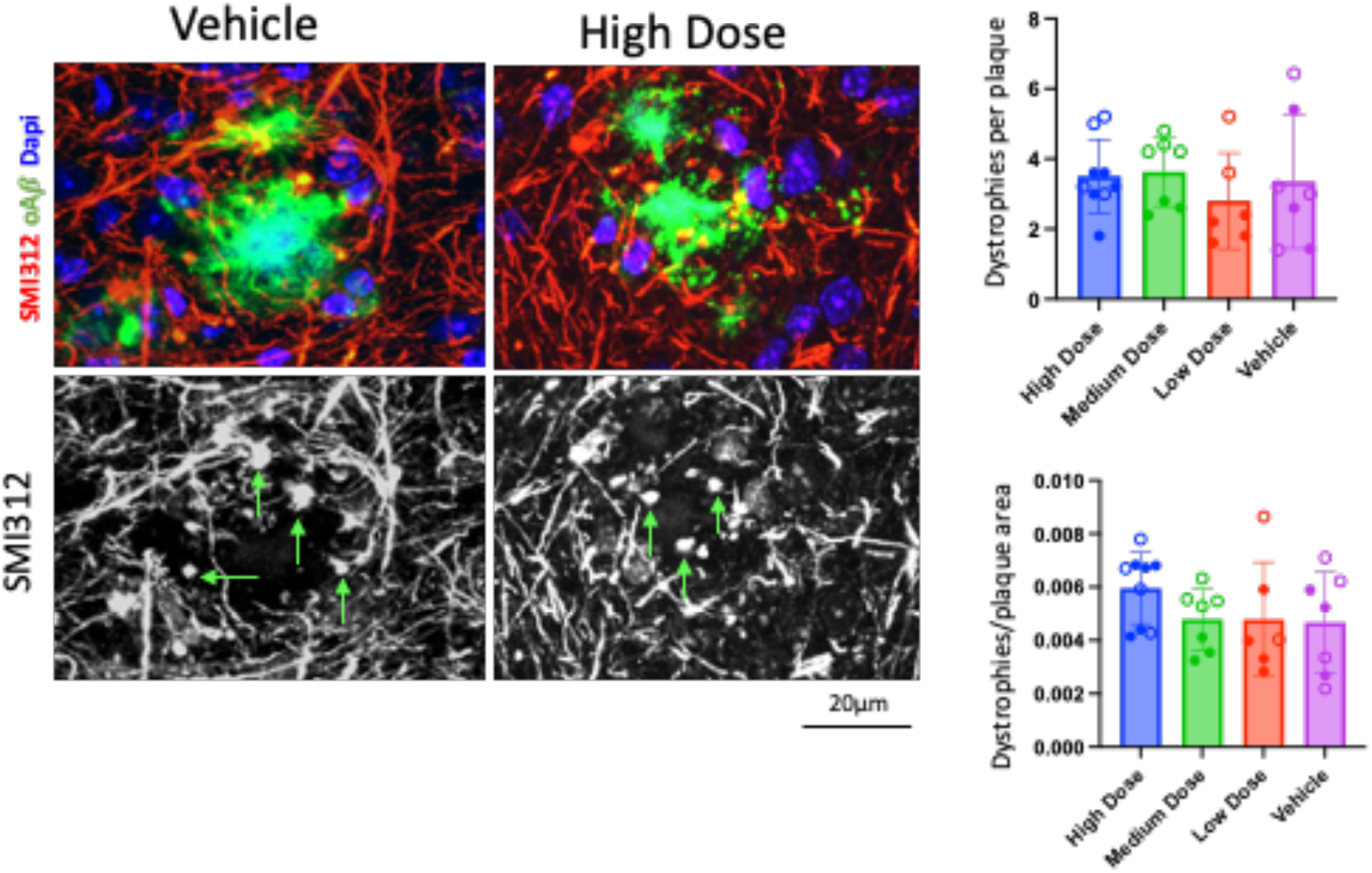
APOE2 does not affect neuritic dystrophies. **(A)** IHC for Smi-312 and Aβ in the cortex of dosed APPPS1/APOE4 animals. **(B)** Analysis shows no difference in the number of dystrophies counted per plaque (F (3, 25) = 0.4710; p=0.7052) even **(C)** when accounting for plaque size (F (3, 25) = 1.127 p=0.3569).

## METHODS

### Study design

Mice were randomly assigned to treatment groups and injected with AAV.APOE2 or a vehicle control 4 months after birth. The impact of human APOE2 on amyloid deposition was assessed 2 months late by IHC and ELISA. Study groups were blinded to the investigator We calculated 8 animals (4 of each sex) per condition would provide a power of > 0.8 for a 30% correction of baseline phenotypes based on prior data ^24^.

### Animals

APPPS1^20^ mice express human mutant APP KM670/671NL and PSEN1 L166P under the Thy1 promotor and have amyloid deposition at 3-4 months of age. This strain was then crossed with humanized APOE4 mouse wherein the mouse *apoE* was replaced with human APOE4 ^19^. Four-month-old APOE4-TR/APPPS1 mice were dosed with high (7E10vg, n=14), medium (2E10vg, n=11), or low (7E9vg, n=11) AAV.APOE2 or a vehicle control (n=10) and followed for 2 months. A cohort of *Apoe* KO (Jackson labs) mice on a C57BL/6 background was included to assess Aβ background levels in tissue and CSF. Experiments were performed in accordance with the National Institutes of Health (NIH) and institutional guidelines and both sexes were used.

Due to the small size of the mouse brain not all animals were used in every analysis and the n indicates numbers used. Open circles indicate females and closed circles indicate males.

### Viral vector construction and production

AAVs were manufactured by the Children’s Hospital of Philadelphia Research Vector Core and resuspended in Diluent Buffer (Research Vector Core). Quality testing and titering was performed in-house by RVC QC and test methods, with procedures and results reported on a Certificate of Analysis (CoA) for each lot.

### Stereotactic intracerebroventricular injections

AAV intracerebroventricular injections were performed as described previously (14, 30). Animals were anesthetized (O_2_/Isoflurane 0.2%) and positioned on a stereotactic frame (David Kopf Instruments). Injections were performed in each lateral ventricle with 5.25 µl of viral preparation using a 33-gauge needle attached to a 10-μl Hamilton syringe (Hamilton Medical) at 0.20 μl/min. Stereotactic coordinates were calculated from bregma (anteroposterior +0.3 mm, mediolateral ±1 mm, and dorsoventral −2 mm).

### Western blot

Mouse cortical tissue was homogenized in 10 volumes by weight of ice-cold TBS with protease and phosphatase inhibitors using a hand-held electric homogenizer. The homogenate was then spun at 10 000g for 10 min and the supernatant (TBS-soluble fraction) was collected for western blot. Protein concentration was determined using a BCA assay. Total protein (5–10 μg) was loaded and separated by 4–12% NuPAGE gels in MES buffer, proteins were then separated by weight for 2 h at 120 V. Proteins were electrotransferred onto nitrocellulose membrane at 30 V for 1.5 h using the XCell II™ Blot Module system in tris-glycine transfer buffer. Membranes were incubated in blocking buffer (Li-Cor Biosciences) diluted 1:1 TBS for 1 h to reduce background staining. Membranes were then incubated with primary antibodies; rb anti-APOE (Novus biologicals, NBP1-31123), and ms anti-GAPDH (Millipore MAB374) diluted in blocking buffer with added 0.1% Tween-20 overnight at room temperature while shaking. Membranes were then washed and incubated with the appropriate 680 and 800 IR dye secondary antibodies (Li-Cor Biosciences). The membranes were imaged using Odyssey infrared imaging system, and analyzed using Odyssey software.

### DNA and RNA extraction and analysis

Genomic DNA was extracted from brain tissue using QIAamp DNA Mini Kit (Qiagen) as per manufacturer’s protocol. Samples were run on BioRad CFX384 Real Time System C1000 Touch using BioRad CFX Manager 3.1 software. Total genome copies were quantified against a 6-point standard curve was generated using linearized plasmid containing the construct. Primer/probes (designed against a non-coding region in the construct) was used with TaqMan® Master Mix (Applied Biosystems).

Total RNA was extracted from brain tissue using TRIzol (Ambion by Life Technologies) as per manufacturer’s protocol. RNA (1 µg) was treated with DNase I, RNase-free (ThermoScientific) as per manufacturer’s protocol. Complementary DNA was generated using the High Capacity cDNA Reverse Transcription Kit (Life Technologies). Samples were run on BioRad CFX384 Real Time System C1000 Touch using BioRad CFX Manager 3.1 software. APOE levels were quantified by designed primer/probes to be used with TaqMan® Master Mix (Applied Biosystems). Exogenous mRNA levels of transgene expression human APOE (Hs00171168_m1) commercial TaqMan® primer/probe set (Applied Biosystems). Endogenous mouse Beta-Actin (Mm02619580_g1) was used as a reference gene to normalize expression across samples.

### ELISA

The concentrations of Aβ40 and Aβ42 were determined by BNT-77/BA-27 (for Aβ40) and BNT-77/BC-05 (for Aβ42) sandwich ELISA (Wako) according to the manufacturer’s instructions.

Aβ40 and Aβ42 concentrations were measured in TBS, SDS-soluble, and SDS-insoluble fractions for each mouse. Sections of mouse brain were homogenized in 10 volumes (w/v) of TBS buffer with a cOmplete protease inhibitor cocktail (Roche), and centrifuged at 1000,000 × g for 30 min at 4 °C. The supernatant was collected and set aside as the TBS-soluble fraction. The pellet was then homogenized in 10 volumes (w/v) of TBS buffer containing 2% SDS, incubated at 37 °C for 30 min and then centrifuged at 100,000× g for 30 min at 20 °C. The SDS-insoluble pellet was dissolved in 500 µl of 70% formic acid and sonicated on ice at 10% power in 1 minute and 30 second intervals until completely dissolved, and then centrifuged at 100,000× g for 30 min at 4 °C. The formic acid-soluble supernatant was desiccated by Speed-Vac and then resuspended 1 volume (w/v) of dimethyl sulfoxide (DMSO). The DMSO-soluble fraction was used as a SDS-insoluble fraction (adapted from Hashimoto et al 2020) ^36^.

### RNAscope

The drop fixed hemisphere of the *Apoe* KO mice was sectioned to 30!m on a freezing ultramicrotome. Three mice per experimental condition (sham injection vs AAV injected) were stained for *APOE* mRNA by RNAscope. RNAscope experiments were performed using the Manual Fluorescent Multiplex kit v2 (Advanced Cell Diagnostics) following manufacturer’s recommendations with minor adjustments. Briefly for each mouse a several sections were baked onto a superfrost slide for use in *APOE* mRNA quantification. Following target retrieval and protease digestion, probe hybridization was carried out at 40°C for 2 h with hs-APOE (433091), 3-plex Positive Control Probe_Mm (320881) and Negative Control Probe-DapB (310043). After amplification steps to obtain the RNAscope signals, the signal was developed using TSA-cy3 (Perkin Elmer FP1170). Sections were counterstained with 1:1000 DAPI, and mounted using immunomount and scanned using an Olympus VS120-S6-W virtual slide microscope, at a magnification of ×10.

To assess microglial reactivity to plaques sections from treated and untreated APOE4/APP/PS1 mice were basked onto a superfrost slide. Following target retrieval and protease digestion, probe hybridization was carried out at 40°C for 2 h with ms-Clec7a-C1 (532061) and ms-P2ry12-C2 (317601-C2), 3-plex Positive Control Probe_Mm (320881) and Negative Control Probe-DapB (310043). After amplification steps to obtain the RNAscope signals, the signal was developed using TSA-cy3 (Fisher Scientific NEL744001KT) for C1 probes and TSA-cy5 (Fisher Scientific NEL745001KT) for C2 probes. Sections were then blocked in 5% NDS in TBS for 1 hour before being incubated with 1:1000 6E10 (BioLegend 803004) overnight at 4°C. Sections were then washed and incubated with Donkey anti-mouse 488 (Invitrogen A-21202) at 1:500 for 1 hour at room temperature. Sections were washed then counterstained with 1:1000 DAPI, and mounted using immunomount and scanned using a nanozoomer (Hamamatsu) at a magnification of 40x Qupath was used to assess the number of RNAscope puncta per cell. An experimenter blind to condition annotated 50 plaques and 50 non-plaque regions per mouse (while only the plaque channel was visible). The plaque annotations were then expanded by 25µm to create the peri-plaque annotation. All annotations were selected for an individual mouse and the positive cell detection tool was used using dapi for the cell nucleus and max nuclear cy5 signal to select microglia without biasing for cells with more or less P2ry12. The subcellular detection tool was then used for both P2ry12 and Clec7a positive puncta with split by shape and intensity selected and with an expected spot size of 0.5µm^2^, a min spot size of 0.25µm^2^, and a max spot size of 3µm^2^.

### Murine IHC

Mice were euthanized by isoflurane inhalation. One cerebral hemisphere was fixed in 4% paraformaldehyde and 15% glycerol in PBS and switched to 30% glycerol in PBS 48 hours later. The remaining hemisphere was snap-frozen for biochemical analysis. Drop fixed hemispheres were processed by neuroscience associates. 40 hemispheres were embedded in a gelatin block and sectioned to 30!m. Sections were permeabilized in 0.5% Triton-X for 15 min before being blocked in 0.1% Triton-X and 5% normal goat serum for 1 h at room temperature. Incubation with primary antibodies was done overnight at 4°C in 0.05% Triton-X and 2.5% normal goat serum. Sections were then washed in TBS and appropriate secondary antibodies were diluted 1:500 in 0.05% Triton-X and 2.5% normal goat serum in TBS at room temperature. Sections were incubated with 1:1000 dapi in TBS for 10 minutes at room temp, washed, and mounted using immunomount.

### Plaque quantification

Every 10^th^ section was stained as described above using rabbit anti Abeta (1:500, IBL, CAT # 18584) for amyloid beta. Amyloid dense core plaques were labeled by 0.05% Thio-S (Sigma-Aldrich) in 50% ethanol before mounting. Sections were mounted and scanned using a nanozoomer microscope at 40x. Sections were quantified using qupath ^37^. For each section cortical areas were selected, and plaques were identified using an object classifier and plaque coverage area was assessed as a percent of the cortical area measured. For plaque size and number, the same sized area was selected in the cortex of each animal and plaques were identified using an object classifier.

### Glial assessment

Several sections were stained as described above. Primary antibodies were 1:1000 biotinylated Ms anti Abeta 82E1 (IBL 10326), 1:500 GFAP-488 (Millipore MAB 3402X), and 1:500 rabbit anti-IBA1 (wako 019-19741). Secondaries were Streptavidin Alexa Fluor 568 (Invitrogen S11226) and Donkey anti-rabbit 647 (A-31573). 5 cortical plaque containing areas were imaged at random from the somatosensory cortex using an Olympus FV3000 confocal laser scanning microscope at 40x. Z stacks were generated and each plaque was assessed on a 4 point scale (Supplemental Figure 2A) for the level of glial reactivity by 2 blind investigators. All plaques for an individual mouse were averaged together to generate the graphs in Figures 3 B and C and Supplemental Figure 2 C and D while all plaques from a given experimental group were assessed for Figure 3D and Supplemental Figure E.

### Synapse quantification

Several sections were stained as described above. Primary antibodies were using 1:500 rabbit anti Abeta 1:500 (IBL, CAT # 18584) and 1:500 goat anti PSD95 (abcam ab12093). Secondaries were Donkey anti-goat 488 (Invitrogen A-11078) and Donkey anti-rabbit 594 (A A-21207). 5 cortical plaque containing areas were imaged at random from the somatosensory cortex using an Olympus FV3000 confocal laser scanning microscope at 60x using an oil diping objective. 5!m of Z stack was imaged at a slice size of 0.56μm. Images were processed using custom Image J and Matlab Macros similar to Jackson et al. ^38^. In brief 10μm x 10μm crops were taken from areas within 15μm of the plaque halo or greater than 40μm from the plaque halo. Cellular debris and dapi were avoided. Crops were thresholded using custom image J macros and synapses quantified using custom matlab macros. All crops were averaged together to find a density near and far from plaques for each mouse.

### Neurite quantification

Several sections were stained as described above. Primary antibodies were using 1:500 rabbit anti Abeta 1:500 (IBL, CAT # 18584) and 1:500 mouse anti SMI312 (Biolegend 837904).

Secondaries were Donkey anti-rabbit 488 (Invitrogen A-21206) and Donkey anti-mouse 594 (A-21203). 5 cortical plaque containing areas were imaged at random from the somatosensory cortex using an Olympus FV3000 confocal laser scanning microscope at 60x using an oil dipping objective. 20μm of Z stack was imaged at a slice size of 1!m. Images were quantified using Image J by a blind experiment who counted number of dystrophies per plaque and also quantified plaque area. In images where more than one plaque was present the largest plaque was quantified.

### APOE2 staining

Several sections were stained as above with notable changes. Primary antibody was 1:200 rabbit anti-APOE2 (Cell Signaling, E7Y30). Primary antibody was incubated at 4°C in 0.05% Triton-X and 2.5% normal goat serum for 1 week to ensure maximum penetration of the tissue. Signal was boosted using a Tyramide SuperBoost kit (Invitrogen, B40925) in with anti-rabbit 594. Sections were imaged using an Olympus FV3000 confocal laser scanning microscope at 40x and Z stacks were generated.

### Human IHC

4µm thick paraffin sections were cut onto a Superfrost slide using a microtome. Following deparaffinization and re-hydration sections were boiled in citrate buffer for 20 minutes before being allowed to cool to room temperature in the citrate buffer. Sections were then washed with TBS and blocked in 10% NDS in TBS for 1 hour at room temperature before being incubated with primary antibody overnight at 4°C. Primary antibodies were 1:500 Iba1 (wako 019-19741) and 1:1000 6E10 (BioLegend 803004) diluted in 5% NDS in TBS. Sections were then washed and incubated with 1:500 Donkey anti-rabbit 594 (Invitrogen A-21207) and 1:500 Donkey anti-mouse 647 (A-31571) in5% NDS for 1 hour at room temperature. Sections were then washed in TBS, before being incubated in 0.05% Thioflavine-S in 50% Ethanol for 8 minutes in the dark. Sections were then dunked in 80% Ethanol 3 times for 10 seconds and then distilled water 3 times for 10 seconds before being counterstained with 1:1000 DAPI, mounted using Immunomount and imaged on an Olympus VS120-S6-W virtual slide microscope, at a magnification of ×20.

### Statistical analyses

Statistical analyses were performed with the Prism software. One-way ANOVA was used to analyze all between group analysis followed by Dunnett’s multiple comparisons test between each group and the vehicle control. Simple linear regression analysis was used to assess the correlation between factors and the number of viral genome copies with p values representing if the slop was significantly non-0. Statistics were performed where each mouse was a single data point. Samples were blinded for each analysis. Human data was assessed as a t-test for microglial reactivity split by APOE4 status and as a simple linear regression where number of APOE4 alleles was treated as an ordinal value with APOE2/3 individuals considered as having -1 APOE4 allele.

### Data availability

The data that support the findings of this study are available from the corresponding author, upon reasonable request

## Acknowledgements

This study was supported by the NIH grants AG047644 (DMH), NS090934 (DMH), T32AG000222-27, P30AG062421, U01NS111671 (BLD, BH), RF1AG047644, the Children’s Hospital of Philadelphia Research Institute and the JPB Foundation.

## Author contributions

RJJ, BLD, and BTH conceived of the idea. RJJ, MSK, JCM, DPF, SED, AM, TN, and LT were responsible for carrying out the experiments. RJJ, MSK, LT, PTR, EC, YC, DMH, BLD, and BTH were responsible for the methodology and resources. RJJ was responsible for writing the original draft and RJJ, MSK, PTR, EC, DMH, BLD, and BTH were responsible for reviewing, editing, and refining the final manuscript. RJJ, BTH, DMH, and BLD acquired the funding that supported this work.

## Declaration of interests

D.M.H. is on the scientific advisory board of C2N diagnostics and has equity. D.M.H. is on the scientific advisory board of Denali Therapeutics, Genentech, and Cajal Therapeutics and consults for Asteroid.

B.L.D. serves an advisory role with equity in Latus Biosciences, Patch Bio, Voyager Therapeutics, Carbon Biosciences, Spirovant Biosciences, Resilience, Panorama Medicines, Saliogen and Homology Medicines. She has sponsored research from Novartis, Roche, Latus, Homology Medicines, Saliogen and Spirovant.

B.T.H. is on the scientific advisory board of Latus Bio and has an equity interest. B.T.H.has a family member who works at Novartis, and owns stock in Novartis; he serves on the SAB of Dewpoint and owns stock. He serves on a scientific advisory board or is a consultant for AbbVie, Aprinoia Therapeutics, Arvinas, Avrobio, Axial, Biogen, BMS, Cure Alz Fund, Cell Signaling, Dewpoint, Eisai, Genentech, Ionis, Latus, Novartis, Sangamo, Sanofi, Seer, Takeda, the US Dept of Justice, Vigil, Voyager.

M.S.K., L.T., Y.H.C., and P.T.R. are founders and shareholders in Latus Biosciences

## References

1. Corder, E.H., Saunders, A.M., Strittmatter, W.J., Schmechel, D.E., Gaskell, P.C., Small, G.W., Roses, A.D., Haines, J.L., and Pericak-Vance, M.A. (1993). Gene dose of apolipoprotein E type 4 allele and the risk of Alzheimer’s disease in late onset families. Science 261, 921–923. 10.1126/science.8346443.

2. Corder, E.H., Saunders, a M., Risch, N.J., Strittmatter, W.J., Schmechel, D.E., Gaskell, P.C., Rimmler, J.B., Locke, P. a, Conneally, P.M., and Schmader, K.E. (1994). Protective effect of apolipoprotein E type 2 allele for late onset Alzheimer disease. Nat. Genet. 7, 180–184. 10.1038/ng0694-180.

3. AlzGene (2017). AlzGene. http://www.alzgene.org/meta.asp?geneID=83.

4. Rebeck, G.W., Reiter, J.S., Strickland, D.K., Hyman, B.T., William Rebeck, G., Reiter, J.S., Strickland, D.K., and Hyman, B.T. (1993). Apolipoprotein E in sporadic Alzheimer’s disease: Allelic variation and receptor interactions. Neuron 11, 575–580. 10.1016/0896-6273(93)90070-8.

5. Reiman, E.M., Arboleda-Velasquez, J.F., Quiroz, Y.T., Huentelman, M.J., Beach, T.G., Caselli, R.J., Chen, Y., Su, Y., Myers, A.J., Hardy, J., et al. (2020). Exceptionally low likelihood of Alzheimer’s dementia in APOE2 homozygotes from a 5,000-person neuropathological study. Nat. Commun. 11, 667. 10.1038/s41467-019-14279-8.

6. West, H.L., William Rebeck, G., and Hyman, B.T. (1994). Frequency of the apolipoprotein E epsilon 2 allele is diminished in sporadic Alzheimer disease. Neurosci. Lett. 175, 46–48. 10.1016/0304-3940(94)91074-X.

7. Serrano-Pozo, A., Das, S., and Hyman, B.T. (2021). APOE and Alzheimer’s disease: advances in genetics, pathophysiology, and therapeutic approaches. Lancet Neurol. 20, 68–80. 10.1016/S1474-4422(20)30412-9.

8. Serrano-Pozo, A., Qian, J., Monsell, S.E., Betensky, R.A., and Hyman, B.T. (2015). APOEε2 is associated with milder clinical and pathological Alzheimer’s disease. Ann. Neurol. 77, 917–929. 10.1002/ana.24369.

9. Castellano, J.M., Kim, J., Stewart, F.R., Jiang, H., B, D., Ronald, Patterson, B.W., Fagan, A.M., Morris, J.C., Mawuenyega, K.G., Cruchaga, C., et al. (2011). Human apoE isoforms differentially regulate brain amyloid-beta peptide clearance. Sci. Transl. Med. 3, 89ra57. 10.1126/scitranslmed.3002156.

10. Deane, R., Sagare, A., Hamm, K., Parisi, M., Lane, S., Finn, M.B., Holtzman, D.M., and Zlokovic, B.V. (2008). apoE isoform – specific disruption of amyloid β peptide clearance from mouse brain. J. Clin. Invest. 118, 4002–4013. 10.1172/JCI36663DS1.

11. Garai, K., Verghese, P.B., Baban, B., Holtzman, D.M., and Frieden, C. (2014). The Binding of Apolipoprotein E to Oligomers and Fibrils of Amyloid-β Alters the Kinetics of Amyloid Aggregation. Biochemistry 53, 6323–6331. 10.1021/bi5008172.

12. Hashimoto, T., Serrano-Pozo, A., Hori, Y., Adams, K.W., Takeda, S., Banerji, A.O., Mitani, A., Joyner, D., Thyssen, D.H., Bacskai, B.J., et al. (2012). Apolipoprotein E, Especially Apolipoprotein E4, Increases the Oligomerization of Amyloid Peptide. J. Neurosci. 32, 15181–15192. 10.1523/JNEUROSCI.1542-12.2012.

13. Fagan, A.M., Watson, M., Parsadanian, M., Bales, K.R., Paul, S.M., and Holtzman, D.M. (2002). Human and murine ApoE markedly alters A beta metabolism before and after plaque formation in a mouse model of Alzheimer’s disease. Neurobiol. Dis. 9, 305–318. 10.1006/nbdi.2002.0483.

14. Shi, Y., Yamada, K., Liddelow, S.A., Smith, S.T., Zhao, L., Luo, W., Tsai, R.M., Spina, S., Grinberg, L.T., Rojas, J.C., et al. (2017). ApoE4 markedly exacerbates tau-mediated neurodegeneration in a mouse model of tauopathy. Nature. 10.1038/nature24016.

15. Klein, R.C., Mace, B.E., Moore, S.D., and Sullivan, P.M. (2010). Progressive loss of synaptic integrity in human apolipoprotein E4 targeted replacement mice and attenuation by apolipoprotein E2. Neuroscience 171, 1265–1272. 10.1016/j.neuroscience.2010.10.027.

16. Koffie, R.M., Hashimoto, T., Tai, H.-C., Kay, K.R., Alberto, S.-P., Joyner, D., Hou, S., Kopeikina, K.J., Frosch, M.P., Lee, V.M., et al. (2012). Apolipoprotein E4 effects in Alzheimer’s disease are mediated by synaptotoxic oligomeric amyloid-β. Brain J. Neurol. 135, 2155–2168. 10.1093/brain/aws127.

17. Serrano-Pozo, A., Li, Z., Noori, A., Nguyen, H.N., Mezlini, A., Li, L., Hudry, E., Jackson, R.J., Hyman, B.T., and Das, S. (2021). Effect of APOE alleles on the glial transcriptome in normal aging and Alzheimer’s disease. Nat. Aging 2021 110 1, 919–931. 10.1038/s43587-021-00123-6.

18. Chernick, D., Ortiz-Valle, S., Jeong, A., Qu, W., and Li, L. (2019). Peripheral versus central nervous system APOE in Alzheimer’s disease: Interplay across the blood-brain barrier. Neurosci. Lett. 708, 134306. 10.1016/j.neulet.2019.134306.

19. Huynh, T.P.V., Wang, C., Tran, A.C., Tabor, G.T., Mahan, T.E., Francis, C.M., Finn, M.B., Spellman, R., Manis, M., Tanzi, R.E., et al. (2019). Lack of hepatic apoE does not influence early Aβ deposition: Observations from a new APOE knock-in model. Mol. Neurodegener. 14. 10.1186/s13024-019-0337-1.

20. Radde, R., Bolmont, T., Kaeser, S.A., Coomaraswamy, J., Lindau, D., Stoltze, L., Calhoun, M.E., Jäggi, F., Wolburg, H., Gengler, S., et al. (2006). Aβ42-driven cerebral amyloidosis in transgenic mice reveals early and robust pathology. EMBO Rep. 7, 940–946. 10.1038/sj.embor.7400784.

21. Carrell, E.M., Chen, Y.H., Ranum, P.T., Coffin, S.L., Singh, L.N., Tecedor, L., Keiser, M.S., Hudry, E., Hyman, B.T., and Davidson, B.L. (2023). VWA3A-derived ependyma promoter drives increased therapeutic protein secretion into the CSF. Mol. Ther. - Nucleic Acids, S2162253123001890. 10.1016/j.omtn.2023.07.016.

22. Chen, W.-T., Lu, A., Craessaerts, K., Pavie, B., Sala Frigerio, C., Corthout, N., Qian, X., Laláková, J., Kühnemund, M., Voytyuk, I., et al. (2020). Spatial Transcriptomics and In Situ Sequencing to Study Alzheimer’s Disease. Cell 182, 976–991.e19. 10.1016/j.cell.2020.06.038.

23. Krasemann, S., Madore, C., Cialic, R., Baufeld, C., Calcagno, N., El Fatimy, R., Beckers, L., O’Loughlin, E., Xu, Y., Fanek, Z., et al. (2017). The TREM2-APOE Pathway Drives the Transcriptional Phenotype of Dysfunctional Microglia in Neurodegenerative Diseases. Immunity 47, 566–581.e9. 10.1016/J.IMMUNI.2017.08.008.

24. Hudry, E., Dashkoff, J., Roe, A.D., Takeda, S., Koffie, R.M., Hashimoto, T., Scheel, M., Spires-Jones, T., Arbel-Ornath, M., Betensky, R., et al. (2013). Gene transfer of human Apoe isoforms results in differential modulation of amyloid deposition and neurotoxicity in mouse brain. Sci. Transl. Med. 5, 212ra161. 10.1126/scitranslmed.3007000.

25. Yamamoto, T., Choi, H.W., and Ryan, R.O. (2008). Apolipoprotein E isoform-specific binding to the low-density lipoprotein receptor. Anal. Biochem. 372, 222–226. 10.1016/j.ab.2007.09.005.

26. Katz, M.L., Tecedor, L., Chen, Y., Williamson, B.G., Lysenko, E., Wininger, F.A., Young, W.M., Johnson, G.C., Whiting, R.E.H., Coates, J.R., et al. (2015). AAV gene transfer delays disease onset in a TPP1-deficient canine model of the late infantile form of Batten disease. Sci. Transl. Med. 7, 313ra180. 10.1126/scitranslmed.aac6191.

27. Liu, G., Martins, I., Wemmie, J.A., Chiorini, J.A., and Davidson, B.L. (2005). Functional Correction of CNS Phenotypes in a Lysosomal Storage Disease Model Using Adeno-Associated Virus Type 4 Vectors. J. Neurosci. 25, 9321–9327. 10.1523/JNEUROSCI.2936-05.2005.

28. Jiménez, A.J., Domínguez-Pinos, M.-D., Guerra, M.M., Fernández-Llebrez, P., and Pérez-Fígares, J.-M. (2014). Structure and function of the ependymal barrier and diseases associated with ependyma disruption. Tissue Barriers 2, e28426. 10.4161/tisb.28426.

29. Efthymiou, A.G., and Goate, A.M. (2017). Late onset Alzheimer’s disease genetics implicates microglial pathways in disease risk. Mol. Neurodegener. 12, 43. 10.1186/s13024-017-0184-x.

30. Hong, S., Beja-Glasser, V.F., Nfonoyim, B.M., Frouin, A., Li, S., Ramakrishnan, S., Merry, K.M., Shi, Q., Rosenthal, A., Barres, B.A., et al. (2016). Complement and microglia mediate early synapse loss in Alzheimer mouse models. Science 352, 712–716. 10.1126/science.aad8373.

31. Leng, F., and Edison, P. (2021). Neuroinflammation and microglial activation in Alzheimer disease: where do we go from here? Nat. Rev. Neurol. 17, 157–172. 10.1038/s41582-020-00435-y.

32. Wang, Y., Ulland, T.K., Ulrich, J.D., Song, W., Tzaferis, J.A., Hole, J.T., Yuan, P., Mahan, T.E., Shi, Y., Gilfillan, S., et al. (2016). TREM2-mediated early microglial response limits diffusion and toxicity of amyloid plaques. J. Exp. Med. 213, 667–675. 10.1084/jem.20151948.

33. Meilandt, W.J., Ngu, H., Gogineni, A., Lalehzadeh, G., Lee, S.-H., Srinivasan, K., Imperio, J., Wu, T., Weber, M., Kruse, A.J., et al. (2020). Trem2 Deletion Reduces Late-Stage Amyloid Plaque Accumulation, Elevates the Aβ42:Aβ40 Ratio, and Exacerbates Axonal Dystrophy and Dendritic Spine Loss in the PS2APP Alzheimer’s Mouse Model. J. Neurosci. 40, 1956–1974. 10.1523/JNEUROSCI.1871-19.2019.

34. Budd Haeberlein, S., Aisen, P.S., Barkhof, F., Chalkias, S., Chen, T., Cohen, S., Dent, G., Hansson, O., Harrison, K., von Hehn, C., et al. (2022). Two Randomized Phase 3 Studies of Aducanumab in Early Alzheimer’s Disease. J. Prev. Alzheimers Dis. 9, 197–210. 10.14283/jpad.2022.30.

35. van Dyck, C.H., Swanson, C.J., Aisen, P., Bateman, R.J., Chen, C., Gee, M., Kanekiyo, M., Li, D., Reyderman, L., Cohen, S., et al. (2023). Lecanemab in Early Alzheimer’s Disease. N. Engl. J. Med. 388, 9–21. 10.1056/NEJMoa2212948.

36. Hashimoto, T., Fujii, D., Naka, Y., Kashiwagi-Hakozaki, M., Matsuo, Y., Matsuura, Y., Wakabayashi, T., and Iwatsubo, T. (2020). Collagenous Alzheimer amyloid plaque component impacts on the compaction of amyloid-β plaques. Acta Neuropathol. Commun. 8, 212. 10.1186/s40478-020-01075-5.

37. Bankhead, P., Loughrey, M.B., Fernández, J.A., Dombrowski, Y., McArt, D.G., Dunne, P.D., McQuaid, S., Gray, R.T., Murray, L.J., Coleman, H.G., et al. (2017). QuPath: Open source software for digital pathology image analysis. Sci. Rep. 7, 16878. 10.1038/s41598-017-17204-5.

38. Jackson, R.J., Rose, J., Tulloch, J., Henstridge, C., Smith, C., and Spires-Jones, T.L. (2019). Clusterin accumulates in synapses in Alzheimer’s disease and is increased in apolipoprotein E4 carriers. Brain Commun. 1, fcz003. 10.1093/braincomms/fcz003.

